# Large Language Models to process, analyze, and synthesize biomedical texts – a scoping review

**DOI:** 10.1101/2024.04.19.588095

**Authors:** Simona Emilova Doneva, Sijing Qin, Beate Sick, Tilia Ellendorff, Jean-Philippe Goldman, Gerold Schneider, Benjamin Victor Ineichen

## Abstract

The advent of large language models (LLMs) such as BERT and, more recently, GPT, is transforming our approach of analyzing and understanding biomedical texts. To stay informed about the latest advancements in this area, there is a need for up-to-date summaries on the role of LLM in Natural Language Processing (NLP) of biomedical texts. Thus, this scoping review aims to provide a detailed overview of the current state of biomedical NLP research and its applications, with a special focus on the evolving role of LLMs. We conducted a systematic search of PubMed, EMBASE, and Google Scholar for studies and conference proceedings published from 2017 to December 19, 2023, that develop or utilize LLMs for NLP tasks in biomedicine. LLMs are being applied to a wide array of tasks in the biomedical field, including knowledge management, text mining, drug discovery, and evidence synthesis. Prominent among these tasks are text classification, relation extraction, and named entity recognition. Although BERT-based models remain prevalent, the use of GPT-based models has substantially increased since 2023.

## INTRODUCTION

Natural Language Processing (NLP), a field at the intersection of computer science, artificial intelligence and linguistics, aims at enabling computers with the ability to comprehend language similarly to how humans do. It has become an indispensable tool for processing and analyzing large volumes of text data, widely adopted in various fields including biomedical research and clinical applications [1]. A pivotal methodological advancement in NLP has been the transformer architecture, which underpins most recent Large Language Models (LLMs), including both BERT, Bidirectional Encoder Representations from Transformers, and GPT, Generative Pre-trained Transformer. The transformer model is appreciated for its efficiency and ability to process data in parallel. This capability, combined with a deep learning architecture that leverages attention mechanisms, allows to handle long dependencies and yields a better contextual understanding [2]. Such advancements have fundamentally altered the landscape of NLP model development. The focus has shifted towards refining and customizing pre-trained language models, moving away from the traditional approach of building models from the ground up. This approach is especially advantageous in specialized fields like biomedicine, where annotated datasets are often limited or difficult to compile. By leveraging the fine-tuning capabilities of LLMs for specific tasks, researchers in the biomedical domain can attain high performance levels, even when working with relatively small datasets [3].

Amidst these advancements, the emergence of generative models marks an evolution towards creative and generative applications, setting a contrast to the discriminative nature of earlier models [4, 5]. While BERT and similar systems excel in task-oriented applications such as sentence classification or entity recognition, generative models are capable of producing coherent and contextually relevant text based on the input they receive. This capability facilitates novel applications such as automated content creation, summarization, and question-answering systems in the biomedical domain. In addition, these models show promising capabilities in exploring instruction tuning and in-context learning. This enables them to comprehend and execute a wide array of tasks using natural language instructions or examples, thus further reducing the reliance on task-specific datasets or extensive fine-tuning [6]. However, generative models also introduce new risks and challenges, such as the generation of plausible but incorrect or misleading information, underscoring the need for careful evaluation and application of these technologies [7, 8].

Given the fast-paced nature of NLP development, there is a need for up-to-date literature overviews. This is particularly true in fields like biomedical research, where the ability to swiftly and accurately process vast amounts of text data can substantially impact patient care and scientific discovery [9]. Thus, this scoping review has three goals: Firstly, we aim to identify and categorize the specific tasks within BioNLP that are currently being addressed using LLMs, e.g., in evidence synthesis, pharmacology, or clinical use cases. Secondly, we aim to map the various LLM architectures employed in these BioNLP tasks. Finally, this study sets out to assess the transparency of methods used in the development of these LLMs, including availability of source code and data as well as used hardware and software.

### Related Work

The work presented in [10] presented the first comprehensive review of the development and landscape of transformer-based biomedical language models. It examined the core implementation principles behind these models and their practical applications for BioNLP tasks with a focus on encoder-based language models. Subsequent work expanded the discussion to include RNN-based models and those that leverage decoders for generative pre-training [3]. Furthermore, the authors extended their exploration of biomedical LLMs to encompass applications beyond textual data, including analyses of proteins, DNA, and text-image pairs. Aligning with our goals, they also provided an in-depth exploration of language model applications within the biomedical domain, offering a thorough classification of techniques, datasets, and benchmarking competitions [3].

Other reviews have concentrated on the use of LLMs within specific niches of biomedical text processing. For instance, how LLMs can enhance downstream tasks in literature-based discovery [11]. Another survey shed light on biomedical and clinical NLP research conducted in languages other than English, focusing on data resources, language models, and common NLP tasks in these languages. This survey also highlighted the growing trend of employing transformer-based language models for various NLP tasks in medical fields [12].

Recent surveys have delved into the unique challenges posed by deploying LLMs in healthcare, especially concerning fairness and accountability. For example, an overview of the technical aspects of development and usage of LLM methods in healthcare, including the specific challenges such as ethical considerations encountered when deploying this technology in the field, has been provided [13]. Similarly, the potential of models such as ChatGPT to enhance medical practices, while addressing concerns about safety, ethics, and maintaining the human element in patient care, has been reviewed in [7]. A related systematic review evaluated the benefits and limitations of ChatGPT in healthcare education, research, and practice, based on an analysis of 60 publications from PubMed/MEDLINE and Google Scholar [14]. Insights into the opportunities and challenges of LLMs for biomedical or clinical usage, including their roles in pre-consultation, diagnosis, data management, and their support for medical education and writing, have also been offered in [15].

In the present scoping review, we conduct an examination of the usage of both discriminative and generative LLM architectures in biomedicine, employing a robust systematic review methodology. We present in-depth quantitative analyses, uncover trends over time, and critically assess transparency and Open Science pracrtices in the BioNLP field, thereby enriching the insights of existing research.

## METHODS

### Study registration

We registered the study protocol on the Open Science Framework (OSF) platform (https://osf.io/bjv24/).

### Literature search

We conducted a search for studies in PubMed and EMBASE, spanning from 2017 (i.e., the inception of transformer-based LLMs [2]) to December 19, 2023. Additionally, we searched Google Scholar for EMNLP and ACL machine learning conference proceedings through the “Publish or Perish” software^1^, employing a search string translated from the one used in our PubMed query. This search string was created by an information specialist at the University Library Zurich. The complete search string can be found in the supplementary data.

### Inclusion and exclusion criteria

We included original research articles that employed LLMs to analyze extensive collections of biomedical texts, e.g., scientific publications, (pre)clinical trial registries, patents, and grey literature. Additionally, systematic reviews and meta-analyses that leveraged LLMs for automating certain tasks like abstract screening were also considered.

We excluded non-English publications as well as text data derived from clinical questionnaires or surveys. Reviews were also excluded but were retained as a supplementary source for additional references.

### Study selection

All retrieved publications were screened using ASReview, which incorporates machine learning algorithms to prioritize relevant studies [16]. We established a rule in advance to cease the screening of abstracts once we encountered thirty consecutive irrelevant abstracts. Following this, any remaining abstracts were excluded from further analysis.

### Data extraction and synthesis

The following data were extracted from available full texts of eligible articles: target NLP application (grouped into pre-specified classes), biomedical domain of the application, target database for the application of the developed methods (e.g., PubMed), data type (e.g., abstracts), data availability, hosted application for end-users, LLM (e.g., BioBERT), programming language (e.g., Python), library/framework (e.g., HuggingFace), reported performance metrics (e.g., F1-score), source code availability, pretraining corpus origin (if pre-training was performed, which data was used, e.g., COVID-19 Open Research Dataset), fine-tuning corpus data/task (e.g., BC5), fine-tuning corpus size, number of tasks/datasets for performance evaluation, hardware type (e.g., GPU). These criteria were developed and iteratively improved over a pilot extraction round conducted by two authors (SED and BVI).

We summarized our findings in a narrative fashion bolstered by descriptive statistics of extracted parameters.

### Risk of bias assessment

We assessed the quality of each study against three predefined criteria: 1) Is the process of splitting training from validation data described? 2) Are there methods described for avoiding overfitting or underfitting? 3) Does the dataset or assessment measure provide a possibility to compare to other tools in the same domain?

### Changes from protocol

While in our initial protocol, we were planning to evaluate all ML-based approaches for BioNLP, we decided to shift the focus to specifically evaluating the emerging field of LLMs.

## RESULTS

### Eligible publications

In total, 13,823 publications were retrieved from our database search. 721 publications were eligible for full-text screening (see supplementary Figure S9). A total of 196 articles were included in our final review, with 137 coming from the biomedical journals and 59 from the NLP conference outputs (Figure 1 **A**). Figure 1 **B** also shows the key sources of these articles, highlighting contributions from acknowledged NLP venues such as the ACL/Findings and the EMNLP conferences, as well as biomedical journals such as BMC Bioinformatics and the Journal of Biomedical Informatics.

**Figure 1.**
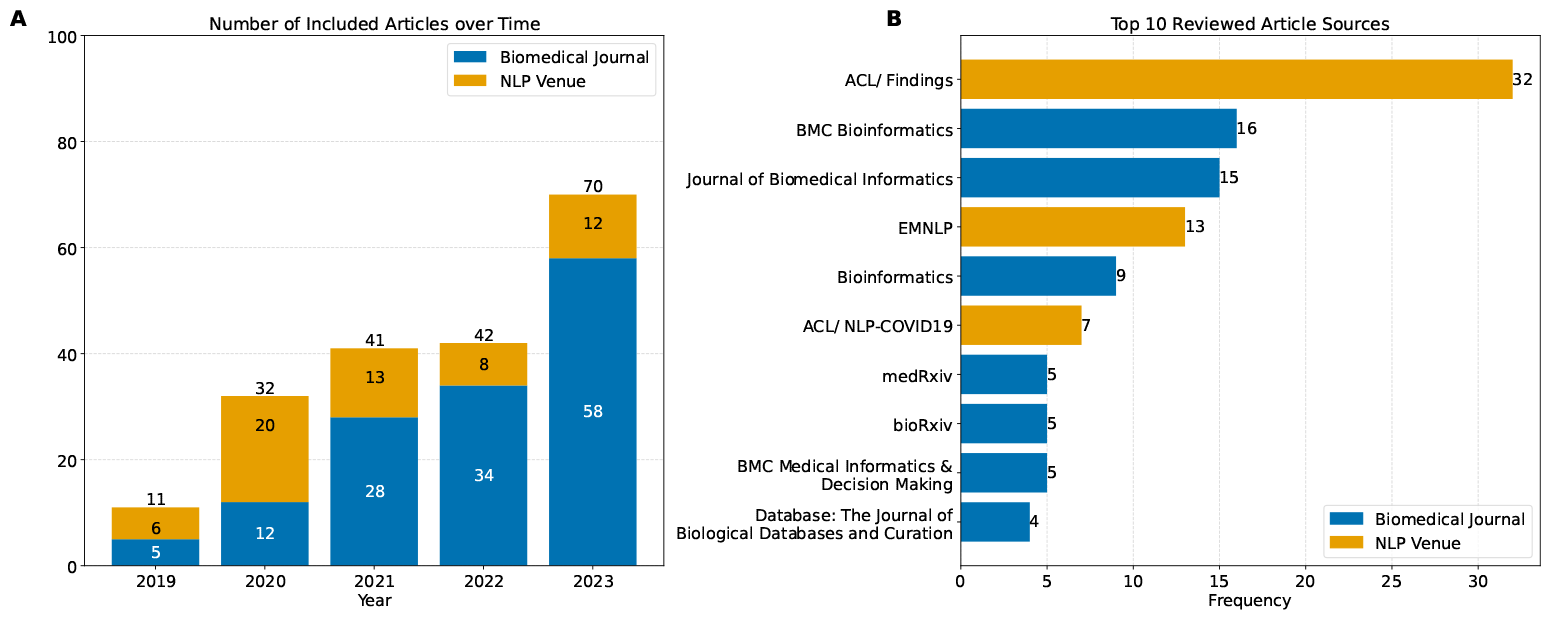
**A**: Number of included articles over the chosen time frame. **B**: Top 10 sources of included articles and their frequency, including NLP venues and biomedical journals.

#### Risk of bias of eligible articles

The overall risk of bias was moderate with 89% of studies reporting whether training and validation data were split, 25% of studies explicitly reported measures to mitigate over-/underfitting of models, and 84% of studies report on benchmarking their approaches against other methods.

### Applications

#### Data Sources and Data Types

Figure 2 introduces the top 10 data sources used to inform the training and refinement phases of the LLMs development. The most frequently reported ones were PubMed/Medline (131 publications, 58%), clinical texts (i.e., textual data generated in clinical settings such as electronic health records; 41, 18%), and social media (16, 7%).

**Figure 2.**
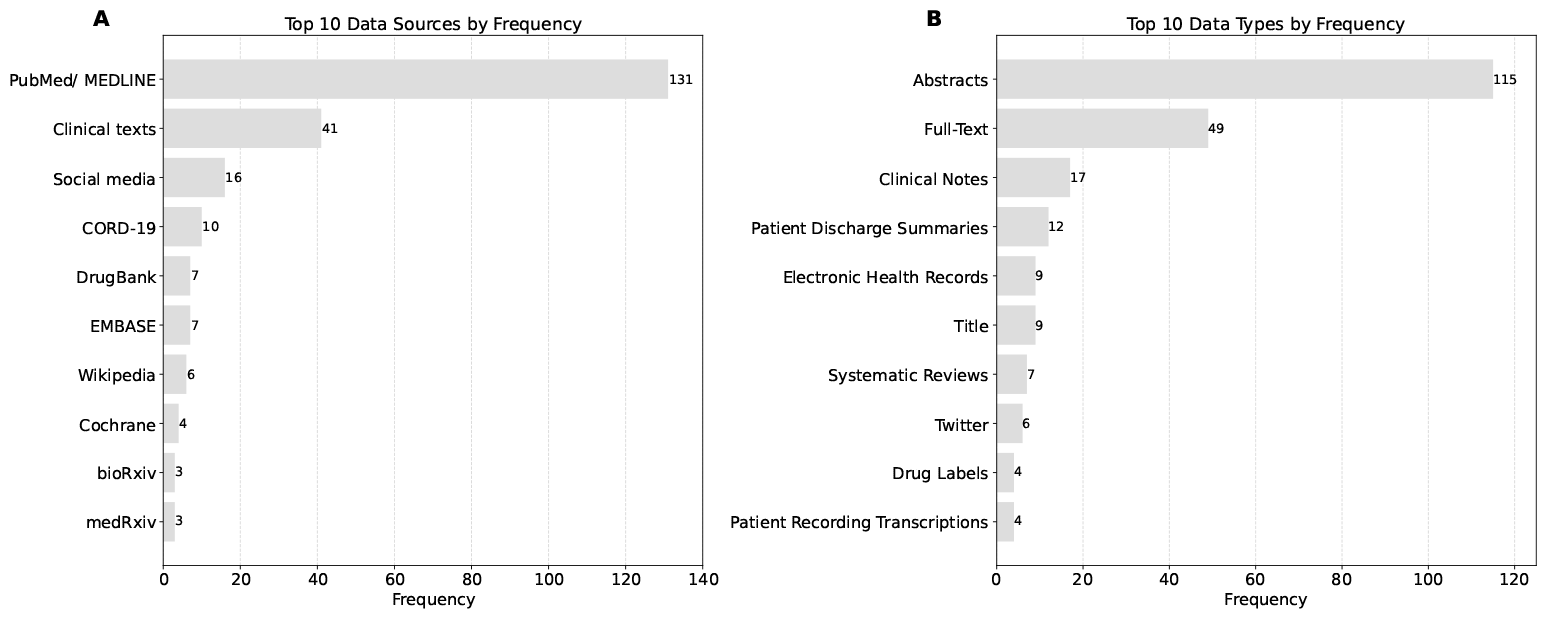
Frequency of specific data sources (**A**) and data types (**B**) used for LLM development and testing. A paper could use more than one data source and type.

The most commonly used data types were abstracts (115 publications, 50%) and full texts of scientific articles (49, 21%) as well as clinical notes (17, 7%) (Figure 2).

#### Biomedical Application Domains

Eligible publications were grouped into 7 main categories of biomedical applications for LLMs (see Table 1). The top three categories were knowledge management (62 publications, 32%), general biomedical text mining (57, 29%), and pharmacology (24, 12%) (Figure 3 **A**). Further details about applications within the top 4 most frequent domains are provided in the supplementary materials in Figure S10.

**Table 1.** Main application domains for LLMs in biomedicine. Abbreviations: NLP, natural language processing.

| Domain Category | Definition |
| --- | --- |
| Knowledge Management | Knowledge management in NLP involves tasks such as literature-based discovery, curation, and screening, focusing on recommendation, summarization, annotation, and categorization of scientific literature. |
| General Biomedical Text Mining | Developing a new/improved methodology for an NLP task (e.g., named entity recognition, dependency parsing). |
| Pharmacology | Development, testing, and understanding of therapeutic agents. |
| Media for Health Care | Analyzing media content to extract health-related information, trends, or public sentiments for healthcare applications. |
| Clinical | Enhancing clinical decision-making, patient care, and medical record management. |
| Evidence Synthesis | The methodological approach used to systematically review, critically appraise, and combine results from multiple studies to draw more robust and comprehensive conclusions about a specific research question or area of interest. |
| Biology | Understanding and categorizing biological processes, mechanisms, and interactions. |

**Figure 3.**
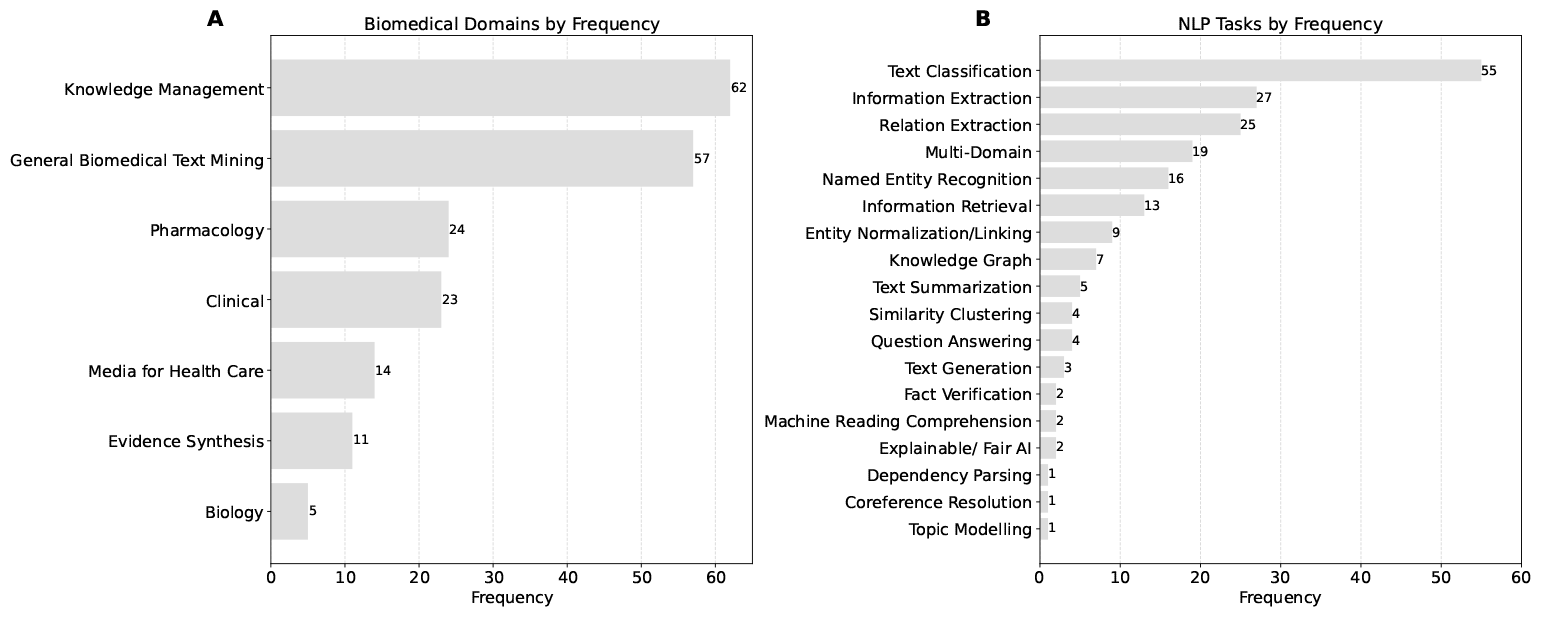
Biomedical domains (**A**) and NLP tasks (**B**) by article frequency. Each paper was assigned to a single primary application domain and one main NLP task.

#### NLP tasks

A variety of NLP tasks has been employed by the eligible articles, most commonly text classification (55 publications, 28%), information extraction (27, 14%), and relation extraction (25, 13%) (Figure 3 **B**). Matching the main domains of application with the NLP tasks shows notable links (Figure 4).

**Figure 4.**
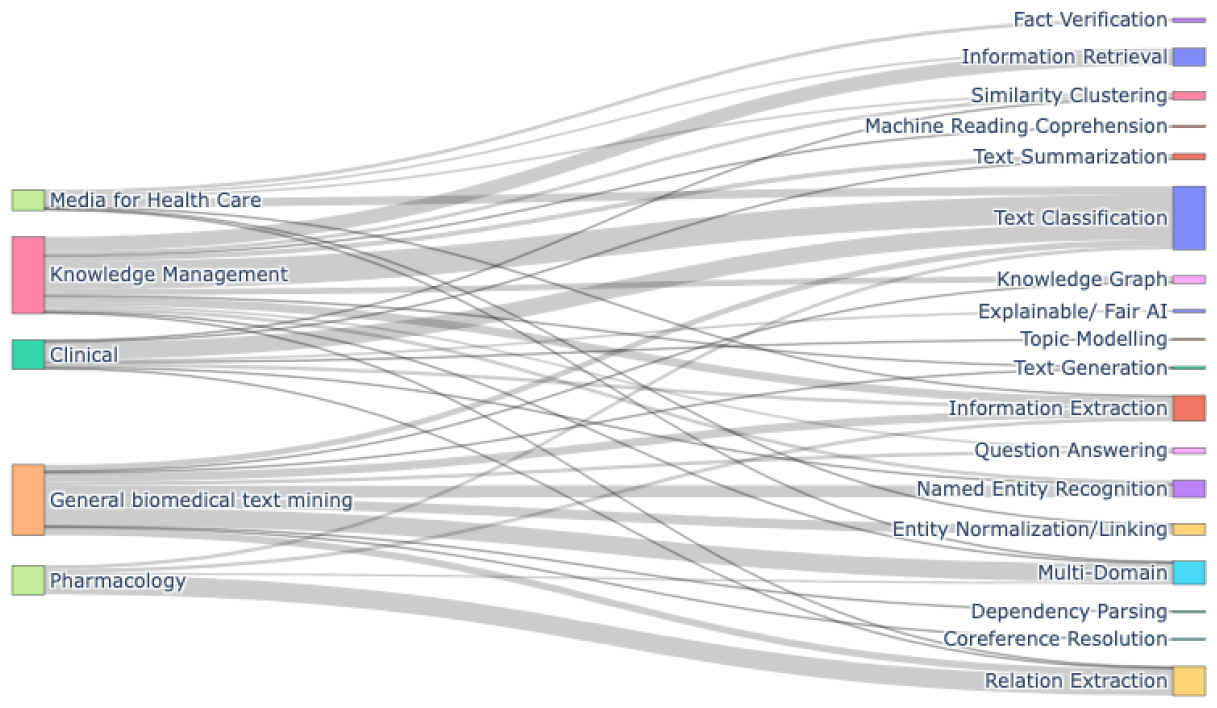
Sankey Diagram showing the relationships between the biomedical domains and the utilized NLP applications. Each domain is represented as a source node, while the associated NLP applications are shown as the target nodes. The thickness of the flows between the two is proportional to the number of articles that exhibit this connection.

In knowledge management, LLMs have been used for two main applications: text classification and information retrieval. Text classification can streamline the organization of large literature volumes by assigning predefined categories or labels to documents. This approach has been effectively applied to enhance reference prioritization in systematic reviews [17, 18, 19, 20, 21]. Text classification has also been used to better understand the structure of papers, for example by automatically predicting sections and headers in Electronic Health Records [22]. Information retrieval focuses on enabling users to find and access the specific information they need from among vast amounts of available content. Various applications have been developed to improve the discovery of novel insights by more effectively aggregating data from existing knowledge databases, as well as improving article recommendation service [23, 24]. For instance, pubmedKB serves as a web server designed to extract and visualize semantic relationships between genes, diseases, chemicals, and variants within PubMed abstracts [25].

General biomedical text mining spans various tasks, with multi-domain analysis standing out. This area focuses on designing methods or frameworks capable of addressing multiple NLP tasks through a unified strategy. An illustration of this is BioGPT, a generative transformer model, which, after being pre-trained on an extensive biomedical literature corpus, demonstrates its versatility across six different BioNLP challenges [26]. Named entity recognition and entity normalization are also challenges commonly tackled together. For example, [27] developed a multi-task model that simultaneously learns sentence-level and token-level labels for NER, utilizing BioBERT for text encoding and sharing hidden states across tasks. [28] introduced an architecture that enhances pre-trained language models by integrating domain-specific knowledge from diverse sources such as the UMLS Metathesaurus and Wikipedia articles on diseases, through knowledge adapters and an attention-based controller, demonstrating performance improvements on various biomedical NLP tasks.

The pharmacological domain utilizes relation extraction the most. This task involves identifying and extracting relationships between entities such as drugs, genes, or proteins from text, which is crucial for pharmacological research and drug development. For example, [29] utilize different BERT models for drug-drug interaction extraction by integrating drug representations from a pharmaceutical knowledge graph and corpus text. [30] used contextual information in extracting long distance adverse drug events from clinical notes with BERT models. There were also overlaps between domains, such as work that introduced a system for detecting adverse drug reactions from text on social media by fine-tuning BERT [31].

In the clinical domain, text classification is frequently utilized for predicting patient outcomes. For instance, in-hospital mortality predictions have been made by combining time series data from various medical devices with clinical notes found in electronic health records [32, 33]. Additionally, there are applications in mental health, such as the automated detection of mental conditions using transcribed patient recordings [34, 35].

Finally, social media platforms have been utilized for mental health screening, employing text classification to identify suicidal risk and predict mental health disorders from user-generated content, such as Reddit and Twitter posts [36, 37, 38]. Additionally, this NLP task has been applied for detecting nuanced emotional states within online health communities [39]. Another recent development has been the use of fact verification techniques to authenticate statements related to COVID-19 [40, 41].

### Large Language Models

#### Models overview and trends over time

The most prominently used LLMs were encoder-based BERT models (275 models, 91%), followed by generative models such as GPT and T5 (27, 9%) (Figure 5).

**Figure 5.**
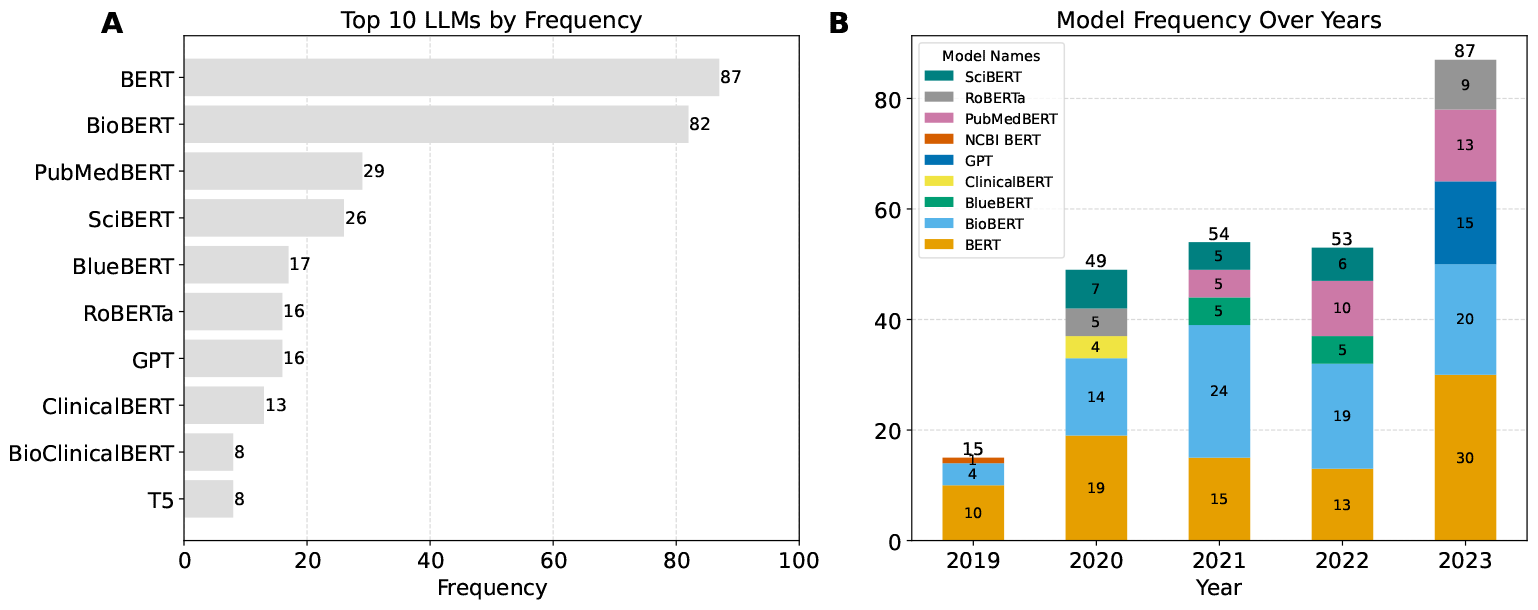
**A**: Most frequently used models. **B**: Top 5 models utilized each year. Each paper could use more than one LLM.

Figure 5 presents the annual usage trends of the top five LLMs. While BERT models are prominently used in earlier years, GPT models see increasing use in 2023. The data also shows that general-purpose models such as BERT are replaced with more specialized models, e.g., BioBERT and SciBERT.

### Technical setup: hardware and programming languages

The most commonly employed hardware were GPUs (92%) (Figure 6). Of note, there was a lack of reporting on hardware use in many instances (118 publications).

**Figure 6.**
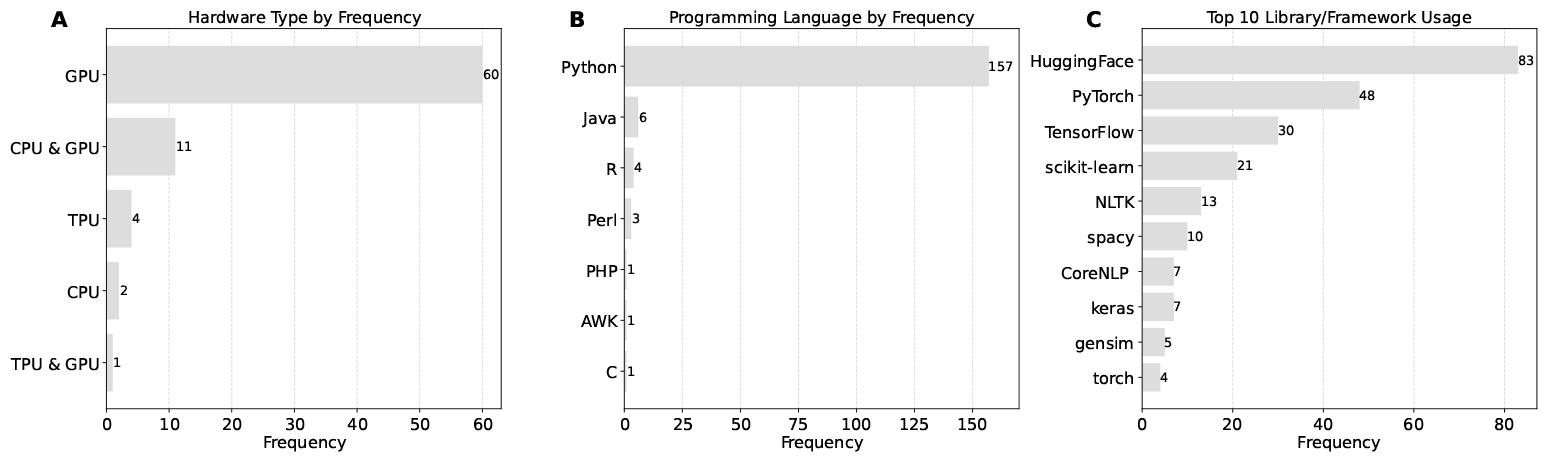
Reported technical setup regarding hardware (**A**), programming language (**B**), and computational library (**C**) used for the method implementation. Each paper could use more than programming language and library.

The vast majority of studies used the Python programming language for their technical setup (157 publications, 91%) (Figure 6). JAVA, R, Perl, PHP, AWK, and C were rarely used. HuggingFace (83 publications, 36%), PyTorch (48, 21%), and TensorFlow (30, 13%) were the most commonly reported computational libraries for using LLMs (Figure 6) . Some studies used more specialized libraries such as Stanford CoreNLP, spaCy, Keras, NLTK, and Torch. Notably, many studies (37) did not report the programming language and computational library used.

### Fine-Tuning Tasks and Datasets

#### Benchmark Datasets

The field of BioNLP relies heavily on the availability of standardized datasets and benchmarks to assess and compare the performance of language models. Table 2 provides a compilation of the most commonly used benchmarks according to the reviewed papers.

**Table 2.** Overview of commonly used datasets for different NLP tasks. The “Frequency” column contains the number of papers that have used the dataset for fine-tuning or testing.

| Task | Dataset | Frequency | Annotation Type | Text Type | Size |
| --- | --- | --- | --- | --- | --- |
| NER | NCBI-disease [42] | 21 | Disease | Abstract | 793 |
|  | BC5-disease [43] | 14 | Disease | Abstract | 1,500 |
|  | BC2GM [44] | 13 | Gene/Protein | Sentence | 20,000 |
|  | BC5-chem [43] | 12 | Chemical | Abstract | 1,500 |
|  | JNLPBA [45] | 9 | Protein, DNA, RNA, cell line | Abstract | 2,404 |
|  | BC4-chem [46] | 6 | Chemical | Abstract | 10,000 |
|  | LINNAEUS [47] | 5 | Species | Full text | 100 |
|  | Species-800 [48] | 5 | Species | Abstract | 800 |
| Relation Extraction | ChemProt [49] | 10 | Chemical-protein interactions | Abstract | 31,784 |
|  | DDIExtraction-2013 [50] | 10 | Drug-drug interaction | Texts and Abstract | 1,017 |
|  | BC5-relations [43] | 9 | Chemical-disease interaction | Abstract | 1,500 |
| Question Answering | BioASQ [51] | 6 | Question-answer pairs and supporting context | Questions with related documents | 4,719 |
| Multiple | MIMIC-III [52] | 8 | Multi-label | EHR | 53,423 |
|  | EU-ADR [53] | 5 | Drug, disorder, targets | Abstract | 100 |
|  | 2010 i2b2/VA [54] | 5 | Multi-label | Clinical reports | 1748 |
|  | Systematic Reviews | 5 | Multi-label | Full text | n.a. |

Many of the reported datasets are collaborative efforts released for competitions and shared tasks, and designed to facilitate fair comparisons among different research teams. Key competitions and shared tasks that emerged from the eligible studies include:

- **Informatics for Integrating Biology and the Bedside (i2b2) Challenges:** The i2b2 Center, active from 2004 to 2014 and funded by the NIH, focuses on releasing clinical records for NLP research. Since 2018, this initiative is officially known as n2c2^1^. The most frequently used dataset from this area is the 2010 i2b2/VA for concepts, assertions, and relation extraction. Other datasets mentioned in the reviewed papers include the 2012 corpus annotated for events, temporal expressions and temporal relations [55], and the de-identification datasets of i2b2 2014 [56]. Three studies utilized data from the 2019 n2c2/OHNLP track on clinical semantic textual similarity [57, 58, 59], and one study tackled the 2022 n2c2 challenge, focusing on medication extraction and event/context classification [60].
- **BioCreative (BC) Workshops:** These workshops host challenges related to the extraction and annotation of biological entities and their relationships from scientific literature. Notable datasets produced from these workshops include BC5, which covers disease and chemical annotations, and BC4, focused on chemical annotations. Additionally, 8 further papers reported utilizing a BioCreative corpus, with LitCovid and DrugProt being the most prominent [61, 62, 29, 63].
- **Text REtrieval Conference (TREC):** TREC is an ongoing series of workshops that cover a wide range of information retrieval topics. Datasets associated with TREC, such as TREC-COVID and Health Misinformation, have been employed in four of the reviewed research papers [64, 65, 66, 67]. One paper also utilized three datasets of the TREC series 2017-2019 for clinical decision support [68].
- **BioNLP Shared Task (BioNLP-ST):** This series is aimed at advancing the extraction of detailed bio-molecular events from scientific texts. The provided data from different years has been utilized nine times in the reviewed papers. For example, the 2011 Protein Coreference dataset is used for the evaluation of methods for coreference extraction among protein/gene [69]. A combination of the datasets from BioNLP 2009, 2011 and 2013 is used in [70] to benchmark a novel fine-tuning approach.

#### Custom-developed Datasets

41% of the publications reported on the development of new datasets. These datasets were mostly in the area of text classification, information retrieval/extraction, and named entity recognition (Figure 7).

**Figure 7.**
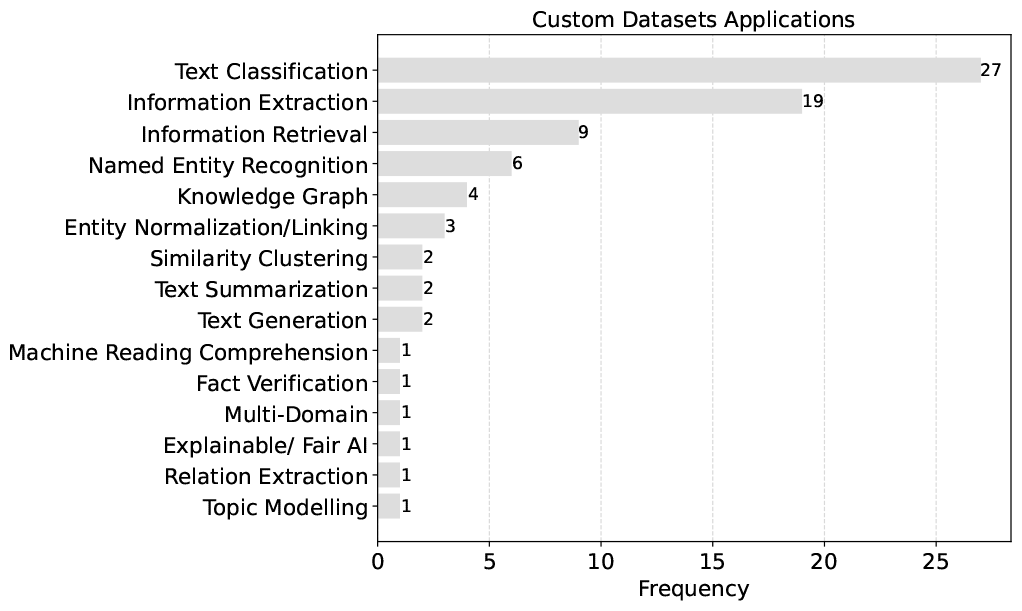
Target applications for which custom data has been annotated or collected.

### Transparency of methods

There was a moderate level of code (e.g., algorithms and computational methods transparency) with fewer than 40% of the studies making their code available alongside the publication (Figure 8). At the same time, there was a high level of data transparency with over three quarters of studies making their datasets fully available. Finally, only a minority of studies hosted applications for end-users with a higher propensity of biomedical journals compared to NLP venues.

**Figure 8.**
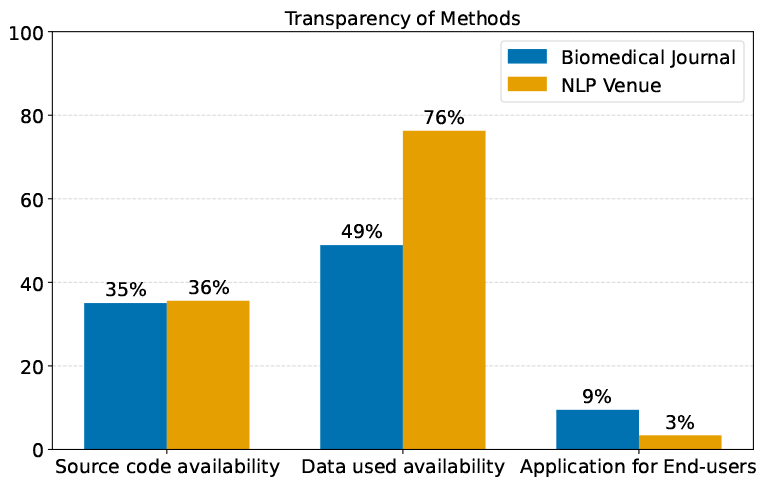
Transparency of LLM Methods: source code and data availability, and end-user application deployment.

## DISCUSSION

### Main Findings

The objective of our scoping review was to explore and organize the specific tasks in BioNLP that are being tackled with LLMs. In the biomedical domain, LLMs have found applications across a diverse set of tasks, including knowledge management, drug discovery, clinical applications, and synthesizing evidence. Within these areas, tasks such as text classification, relation extraction, and named entity recognition/information extraction stand out for their prominence. The primary sources of data for these studies are often PubMed/Medline abstracts or clinical documents, such as electronic health records. While BERT-based models continue to be widely used, there has been a noticeable surge in the adoption of GPT-based models since 2023. Moreover, the technical frameworks for these studies typically involve the Python programming language, with HuggingFace and PyTorch being the most popular computational libraries.

### Limitations and Challanges of LLMs for BioNLP

Most LLM applications reviewed in the present study were developed using information from scientific abstracts. This preference might be due to the easy access to and the concise nature of this data type. However, information from abstracts might not be well aligned with the corresponding full reports. For example, [71] analyse the state of reporting of primary biomedical research and find inconsistencies with respect to the reported sample sizes, outcome measures, result presentation and interpretation, and conclusions or recommendations between abstracts and full texts. This suggests the need to be cautious when developing applications that rely only on the information reported in abstracts, especially for evidence-synthesis. Article abstracts and article bodies can differ not only in content, but also in their structural, linguistic, and semantic composition [72]. LLMs are generally well equipped to handle a wide variety of linguistic styles and structures due to their vast training on diverse text corpora. However, those models can still struggle with the subtlety and specificity required in academic texts. This limitation can be mitigated by the usage of domain specific language models, that also include full-text academic articles in their training data [73].

The most common text-based applications of LLMs in the biomedical field were within the knowledge management domain, showcasing their potential as a solution to the challenges brought on by the exponential growth of biomedical literature [74, 75, 76, 77, 78]. However, we also observed that LLMs have been mostly used as a supportive tool by assisting with preliminary tasks in biomedical research, but they have not yet been capable of replacing more nuanced and critical tasks entirely. For example, while LLMs can expedite the initial screening of literature for relevant studies and assist in the extraction of data from texts, they cannot replace the expert analysis required for interpreting study results, assessing the quality of evidence, or making clinical decisions [79, 80].

Further considerations include the challenge of maintaining model accuracy amidst the fast pace of medical knowledge evolution, the models’ limited interpretability, susceptibility to hallucinations and biases, and ethical and privacy concerns related to handling sensitive patient information. These issues emphasize the importance of cautious and responsible LLM implementation in the biomedical domain [8, 81, 7, 82, 83].

### Emergence of generative LLMs

While BERT-based models have been the predominantly used LLMs, there was a notable surge in GPT-like models in 2023. Generative models have been benchmarked against BERT-based approaches as a classification and information extraction technique for biomedical literature curation [84, 85, 86]. Notably, they have also been applied in innovative ways, such as generating symptom definitions to enrich datasets and improve annotator precision, as well as for summarizing radiology reports, potentially aiding clinical decision-making [87, 88]. ChatGPT’s interactive capabilities have also been explored for preliminary screening in cases of mild cognitive impairment [89].

Recently, the landscape of generative biomedical models has been enriched by open-source solutions such as LLama 2, MEDITRON, and BioMistral [90, 91, 92]. These models are rapidly narrowing the performance gap with proprietary counterparts, while also promoting a more transparent, reproducible, and independent research ecosystem. Furthermore, the development of streamlined versions, such as ollama, optimized for local computing resources, highlights this trend by enhancing the practicality and accessibility of LLMs for a broader range of research applications^1^.

### Transparency of methods

Finally, while a minority of studies present end-user applications, a notable portion leverages openly available datasets or share their proprietary datasets. Despite this positive trend, only around one third of these studies includes references to their code repositories, a critical factor for ensuring the validity, reproducibility, and generalizability of their methods [93, 94]. While guidelines for reporting experimental setups and results exist the NLP and machine learning fields, there seems to be a need for further approaches to improve the reporting rates [95, 96, 97]. Moreover, the use of API-based models such as GPT introduces additional challenges for research transparency and reproducibility [98]. Shifting towards open-source models can address these issues by providing full access to the models’ codebase, fostering an environment where research findings can be easily replicated, scrutinized, and extended by the wider scientific community.

## LIMITATIONS

First, the current literature review is primarily centered on text-based applications of LLMs in biomedicine which stands in contrast to the multimodal data of biomedical information. However, we believe that textual data is still an underused data type to inform the biomedical field.

Second, we omitted articles published in languages other than English, which might restrict the scope of this review.

## CONCLUSION

This scoping review provides an overview how LLMs are used for NLP tasks in biomedicine and health sciences. The findings from our scoping review underline the rapid progress of LLMs, emphasizing their potential in accelerating discovery and enhancing health outcomes. These advancements signal a promising avenue for leveraging LLMs in biomedicine and health, given their capacity to process and analyze complex biomedical texts with high proficiency. However, we also acknowledges the inherent risks and challenges associated with the deployment of LLMs such as ChatGPT in these sensitive areas, including evidence synthesis. Issues such as the generation of fabricated information and concerns over legal and privacy implications pose significant hurdles to the safe and ethical application of these technologies. These challenges highlight the need for careful consideration and management to mitigate risks associated with the use of LLMs in biomedicine and health.

## DECLARATIONS

### Acknowledgments

We thank Alisa Berger for conducting the comprehensive literature search.

## Data and code availability statement

All data and code that support the findings of this study are available in a public GitHub repository^1^. For any questions regarding the data, meta-data, or analysis code, contact the corresponding author, SED.

## Author contributions

Conception and design of study: SED, BS, BVI; Acquisition of data: SED, SQ, BVI; Analysis of data: SED; Interpretation of data: SED, SQ, BSn TE, J-PG, GS, BVI; Drafting the initial manuscript: SED; All authors critically revised the paper draft.

## APPENDIX

### Search string

Supplementary search string (‘data mining’/exp OR ((data NEXT/1 mining):ti,ab,kw)) AND (literature:ti,ab OR abstract:ti,ab OR abstracts:ti,ab OR text:ti,ab OR articles:ti,ab) OR (((analyz* OR analys* OR extract* OR screen* OR evaluat* OR classif* OR ‘natural language processing’) NEAR/3 (literature OR abstract OR abstracts OR text OR articles)):ti,ab,kw) OR ((text NEXT/1 mining):ti,ab,kw) AND ‘natural language processing’/exp OR ‘artificial neural network’/exp OR ‘support vector machine’/exp OR ‘machine learning’/de OR ‘automated pattern recognition’/exp OR ‘artificial intelligence’/de OR ‘semi automation’:ti,ab,kw OR automation:ti,ab,kw OR ‘artificial intelligence’:ti,ab,kw OR ai:ti,ab,kw OR ‘natural language processing’:ti,ab,kw OR ((machine NEXT/1 (intelligence OR learning)):ti,ab,kw) OR (((‘text mining’ OR ‘data-mining’) NEXT/3 (tool* OR technique* OR system)):ti,ab,kw) OR (((deep OR comput* OR model* OR convolutional OR artificial OR algorithmic OR connectionist OR mathematical) NEXT/1 ‘neural network*’):ti,ab,kw) OR ((ann NEXT/3 (analysis OR approach OR method* OR model* OR output OR technique* OR training*)):ti,ab,kw) OR ((connectionist NEXT/1 (model OR network)):ti,ab,kw) OR ((‘support vector’ NEXT/1 (machine* OR classif* OR network OR regression)):ti,ab,kw) OR ((automated NEXT/3 (tool* OR technique* OR system OR ‘pattern recognition’ OR analyz* OR analys* OR extract* OR screen* OR evaluat* OR classif*)):ti,ab,kw)

### Flow diagram of the excluded and included studies

Figure 9 illustrates the process of the scoping review, showcasing the flow of information through the different phases of a literature review, including identification, screening, eligibility, and inclusion of studies.

**Figure 9.**
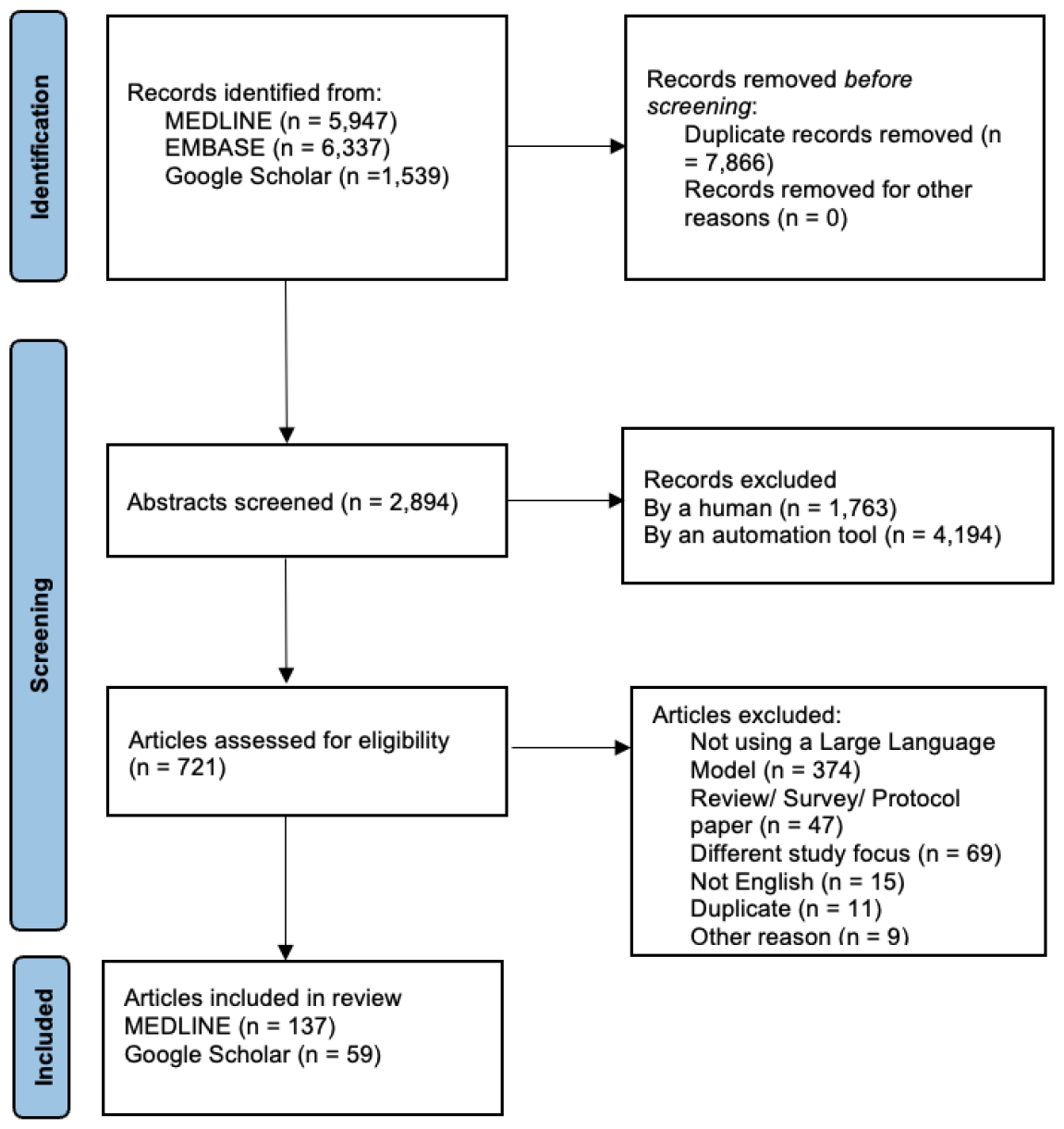
PRISMA Flow Diagram [99].

### Target Biomedical Applications Subdomains

Further details about downstream applications within the top 4 most represented biomedical domains are provided in Figure 10. These subcategories have been identified based on the objectives outlined in the reviewed papers.

**Figure 10.**
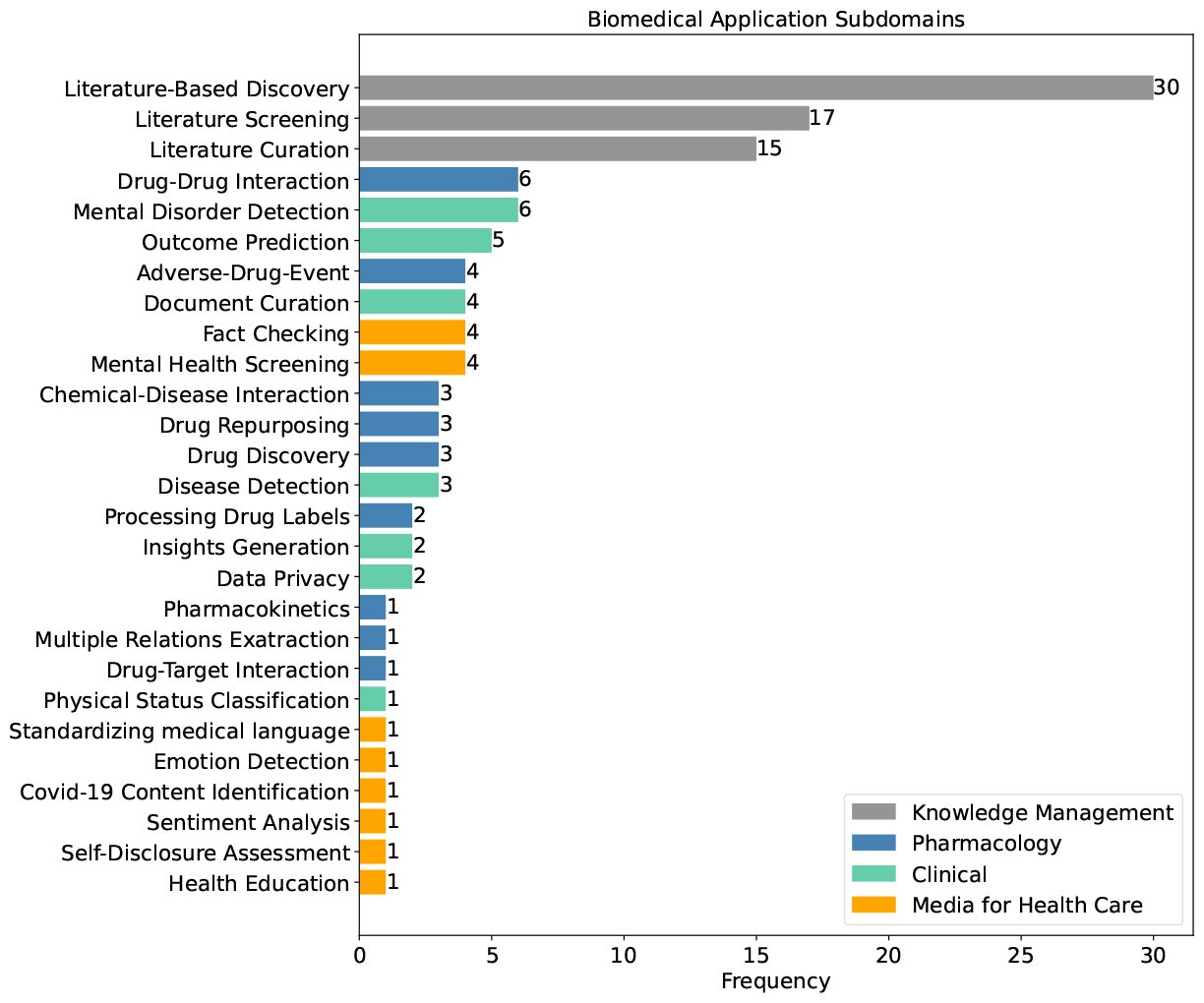
Number of articles assigned to each subdomain.

### Number of tested models per paper over time

The mean number of LLMs used by individual papers ranges between 1.36 and 2.39, with a slight upward trend over time (Figure 11).

**Figure 11.**
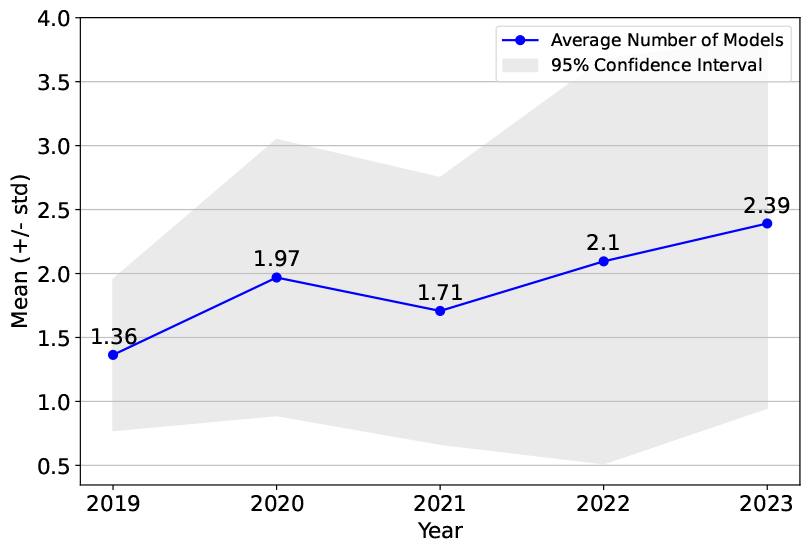
Average number of different models used per paper each year.

### High-level LLM Background

The GPT-like and BERT-like models represent two distinct LLM approaches, differentiated by their architecture, training methods, and use cases [100].

### BERT-style Language Models

BERT employs a bidirectional framework, analyzing text in both directions simultaneously, which is enabled by its use of the Transformer encoder architecture. This bidirectionality allows BERT to understand the context of a word based on its entire surrounding text, making it good at tasks that require a deep understanding of language context, such as sentiment analysis, question answering, and named entity recognition. BERT’s pre-training involves masked language modeling and next sentence prediction, tasks that help it learn a comprehensive understanding of language structure and flow.

While most reported models follow a BERT-based architecture, they differ in the pretraining corpus used. The standard BERT [101] model is pretrained on texts from Wikipedia and BookCorpus, which is considered general-domain. To improve its performance in BioNLP tasks, the models can be trained on biomedical text corpus in two ways [102]:

1. Mixed-Domain Pretraining: Starting with a general-domain BERT model training is continued using biomedical texts. In the case of BioBERT [103] this includes PubMed abstracts and PubMed Central (PMC) full-text articles. BlueBERT [104] uses both PubMed abstracts and de-identified clinical notes from MIMIC-III [52].
2. Domain-Specific Pretraining: In this setup the language model is trained using purely in-domain data. PubMedBERT [102] follows this approach and is pretrained from scratch on abstracts and full-texts from PubMed only.

SciBERT [105] is also trained from scratch on a purely scientific corpus from Semanitic Scholar. However its pretraining corpus is a mixture of biomedical and computer science texts. An overview of the most frequently used BERT-based models in the reviewed literature is given in Table 3.

**Table 3.** Training corpora for common BERT-based models.

| Training Corpus | BERT | BioBERT | PubMedBERT | SciBERT | BlueBERT |
| --- | --- | --- | --- | --- | --- |
| General | ✓ | ✓ | ✗ | ✗ | ✓ |
| PMC | ✗ | ✓ | ✓ | ✗ | ✗ |
| PubMed | ✗ | ✓ | ✓ | ✗ | ✓ |
| Semantic Scholar | ✗ | ✗ | ✗ | ✓ | ✗ |
| Clinical Notes | ✗ | ✗ | ✗ | ✗ | ✓ |

#### GPT-style Language Models

GPT-like models operate on a unidirectional framework, processing text from left to right and utilizing the Transformer decoder architecture, making it well fit for generative tasks such as text completion and content creation. Its training is focused on predicting the next word in a sequence, optimizing the model for generating coherent and contextually relevant text. While BERT-like architectures typically require task-specific fine-tuning to achieve optimal performance, models like GPT-3 demonstrate few-shot and zero-shot learning capabilities. Few-shot learning refers to the model’s ability to perform tasks with a very limited amount of training data, while zero-shot learning refers to its ability to perform tasks without any task-specific training data. This is achieved through advanced prompting techniques and in-context learning, where the model generates responses based on the context provided in the prompt [106].

## Footnotes

1 https://harzing.com/resources/publish-or-perish

3 https://portal.dbmi.hms.harvard.edu/projects/n2c2-nlp/

3 https://ollama.com/

4 https://github.com/Ineichen-Group/LiteratureReview-LLM-Biomedicine/tree/main

